# Lessons from the equator: a window on the future of vibriosis in a warming Earth

**DOI:** 10.64898/2026.03.03.709458

**Authors:** Christopher Colin Neoh, Eric Dubois Hill, Fabini D. Orata, Craig Baker-Austin, Marisa Hast, Michael J. Hughes, Bryan M.H. Keng, Patrick Martin, Tan Yen Ee, Crystal Shie Lyeen Wong, Chew Ka Lip, Su Gin Douglas Chan, Chen Yihui, Timothy Barkham, Swapnil Mishra, Jaime Martinez-Urtaza, Alison E. Mather, Christine Lee, Yann Felix Boucher

## Abstract

Vibrios are important aquatic human pathogens, causing gastroenteritis, wound infections, and cholera. Vibriosis (non-cholera infection) has been increasing in temperate climates due to correlation between *Vibrio* abundance and water temperature. However, how warming oceans differentially influence various *Vibrio* species, and whether incidence is higher in tropical regions, remains understudied. We assembled the first comprehensive dataset of culture-confirmed vibriosis cases outside a temperate climate from equatorial Singapore. Comparison to cases from the detailed United States (US) CDC COVIS system revealed that vibriosis incidence is over two times higher in Singapore than the United States. Causative species also differ markedly, with *Vibrio fluvialis* and non-toxigenic *Vibrio cholerae* dominating in Singapore (>40%) but are much less prominent in the United States (<15%). Singapore cases have remained stable in the last decade, while US incidence is rising rapidly, especially for *V. fluvialis* and *V. cholerae* (13% and 17% per year). Differences in causative species in Singapore and the United States suggest ecological factors may shape infection dynamics, with some species benefitting more from warming oceans, raising their equatorial prominence. Incidence is stable in Singapore, likely due to slower warming of waters near the equator. In contrast, vibriosis in the United States will likely continue to increase steadily, with changing species composition, likely becoming tropicalized.

## INTRODUCTION

Members of the genus *Vibrio* are the most important microbial pathogens from marine environments and coastal waters, where they can cause an array of human infections.^1^ Only a single lineage from the *Vibrio cholerae* species is responsible for the most notorious disease in this group, cholera, caused by pandemic (toxigenic) strains.^2^ However, the genus includes a diversity of other pathogenic species of global concern – *Vibrio vulnificus, Vibrio parahaemolyticus, Vibrio fluvialis*, and *Vibrio alginolyticus*. Infections caused by these, along with those from non-toxigenic *V. cholerae* lineages, are collectively termed vibriosis.^3^ Many of these bacteria can cause sporadic but potentially severe gastrointestinal illnesses and wound infections, which can advance to severe outcomes such as necrotizing fasciitis, amputation, septicemia, and death. Several factors common to these bacteria, including their environmental ubiquity, sensitivity to temperature, rapid replication capabilities, and an unusual genetic structure coupled to widespread genomic rearrangement make them formidable emerging (and re-emerging) pathogens.^3^ The close relationship between climatic anomalies and reported incidents associated with non-cholera vibrios have caused concern, with infections increasingly reported in colder/temperate regions. The relationship between climate and incidence of vibriosis is well established and the number of infections they cause has increased dramatically over the last two decades.^1^ In coastal areas with salinity optimal for their growth, the environmental abundance of vibrios correlates tightly with water temperature.^1,4^ In temperate countries, *Vibrio* infections follow seasonal patterns, with higher incidence in warmer months.^5^

Most of these insights have come from the Cholera and Other Vibrio Illness Surveillance (COVIS) database of the US Centers for Disease Control and Prevention (CDC), which began nationwide mandatory reporting of all *Vibrio* infections in 2007.^6^ COVIS remains the most comprehensive vibriosis database globally. However, the paucity of data from other countries, especially those theoretically more vulnerable to such infections due to warmer climates, has limited our global understanding of vibriosis. Differences between *Vibrio* species in terms of their ecology and clinical profile are insufficiently understood, as many epidemiological analyses tend to aggregate all *Vibrio* species into a single group. Since species vary in their environmental niches, modes of transmission, and disease outcomes, genus-level aggregation can mask important patterns.^3^ Species-resolved surveillance and analyses are therefore essential for capturing species-specific trends and disease dynamics.

In this study, we address these gaps by analyzing detailed infection data from the past decade in Singapore, a country in Southeast Asia near the equator, and comparing it with incidence patterns of all major pathogenic *Vibrio* species across different regions of the United States since 2007.

## RESULTS

### Geographical variation of vibriosis in the United States

In the United States, the causative agents of vibriosis infections showed strong geographical variations. Based on similarities in the relative proportion of vibriosis cases attributed to each *Vibrio* species, six broad regional groups were identified (Fig. 1 and Supplementary Fig. 1). Among Pacific states, species profiles differed by latitude. In Northern Pacific states, most infections (89%) were caused by *V. parahaemolyticus*. These states reported high incidences for this species, ranking among the top four in the country (Hawaii being the third highest overall) (Fig. 2 and Supplementary Fig. 2). The Southern Pacific (California) had a lower relative proportion of *V. parahaemolyticus* infections compared to its northern neighbors. The odds of a vibriosis case being caused by *V. parahaemolyticus* were 3.6 times higher in the Northern Pacific than in the Southern Pacific (OR = 3.60; SE = 0.65; 95% CI = 2.41–5.38; *z* = 7.14; *p* < 0.001).

**Fig. 1.**
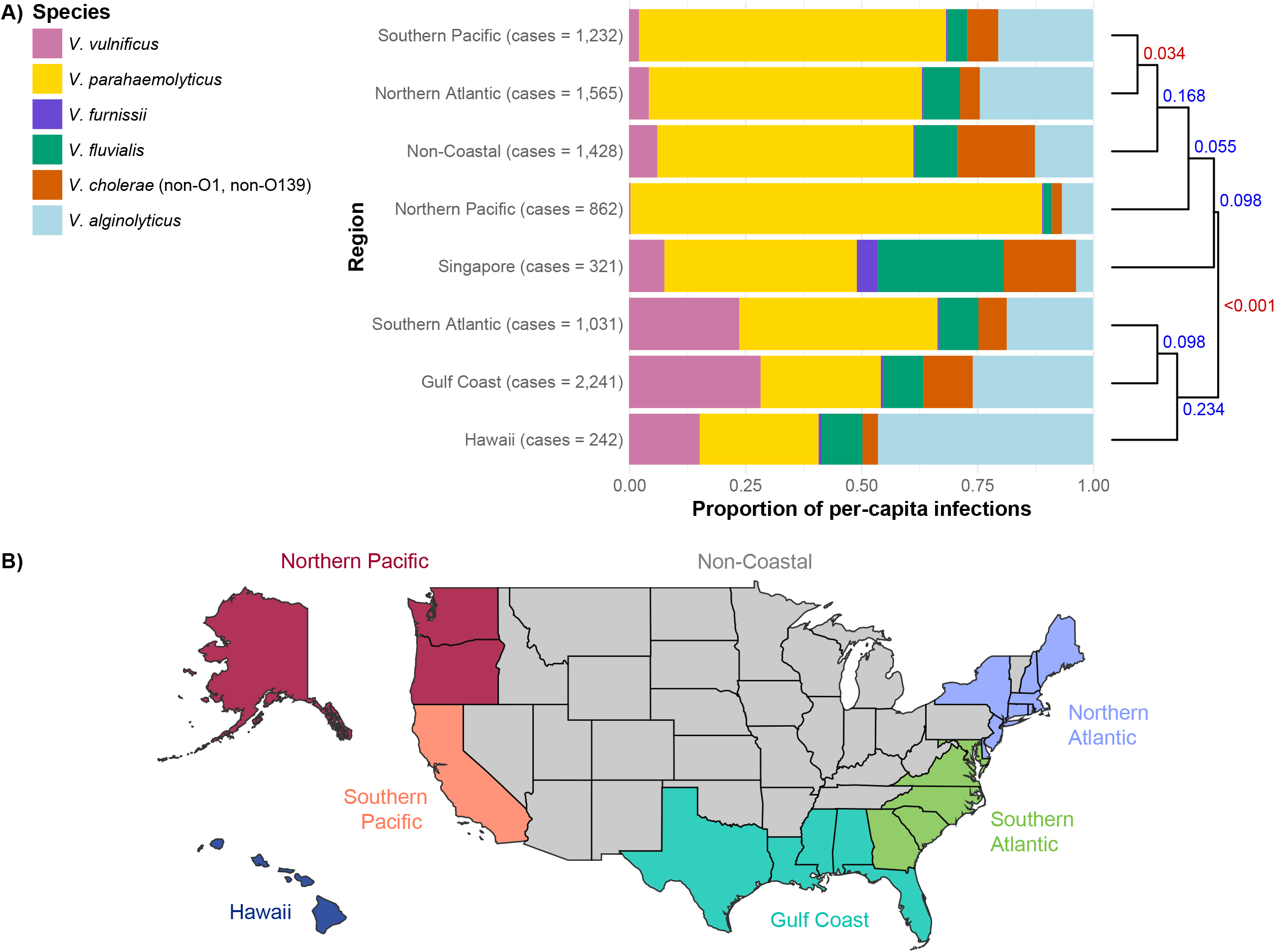
Relative proportion of *Vibrio* infections by causative species across regions of the United States and Singapore. (A) Regions were grouped with agglomerative hierarchical clustering based on Bray–Curtis dissimilarities of their species-level relative-abundance profiles, using the average-linkage (UPGMA) method to merge clusters. *p*-values test the null hypothesis that the cluster does not exist, with red (*p* ≤ 0.05) denoting statistically significant clusters and blue (*p* > 0.05) indicating non-significant groupings. (B) Map of the United States highlighting the regional groupings used in the analysis, which were determined by hierarchical clustering at the state level (see Supplementary Fig. 1).

**Fig. 2.**
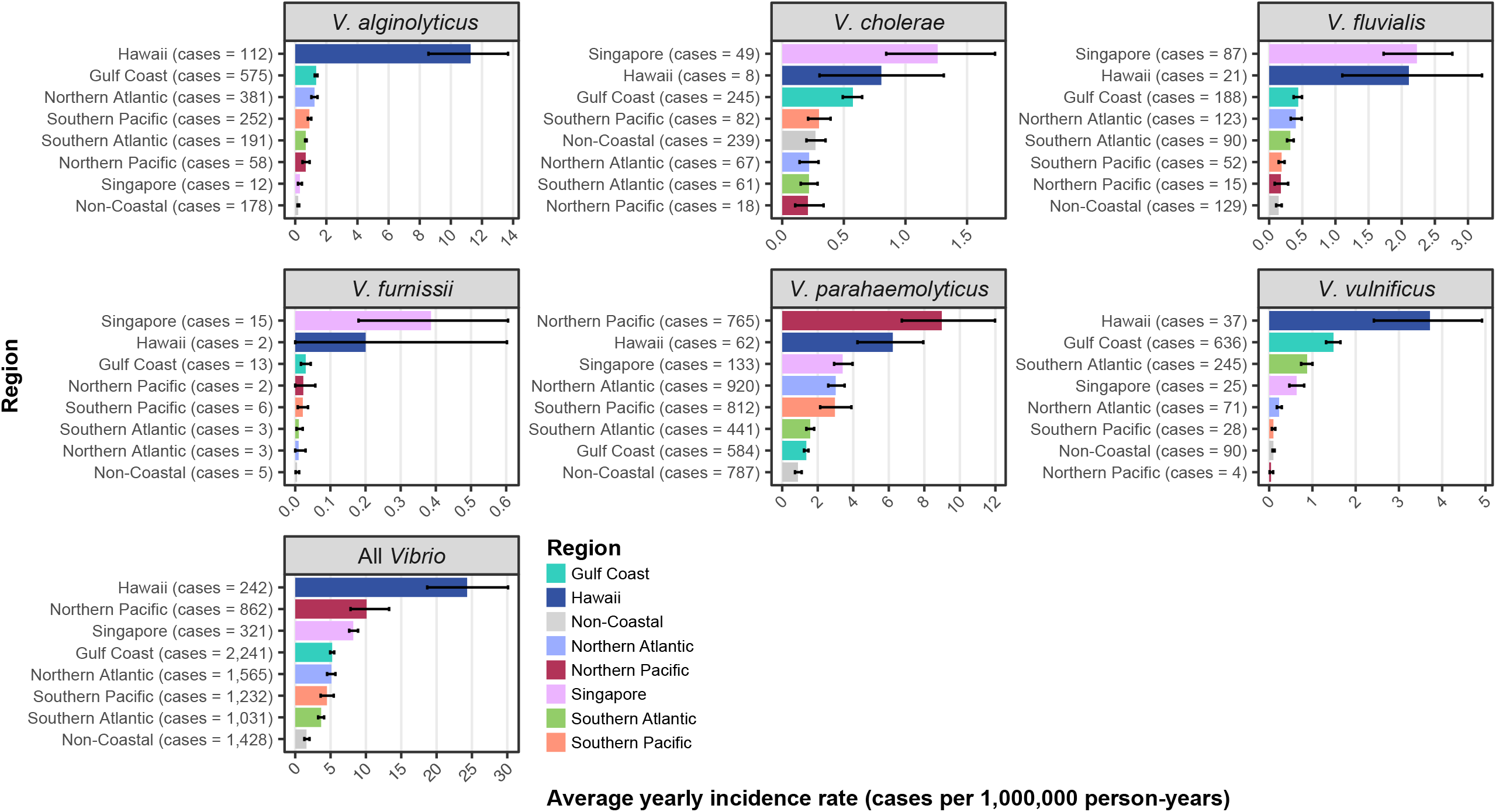
Incidence rates of vibriosis by causative species across regions of the United States and Singapore. Incidence rates were calculated for 2013–2019 from the US COVIS database and records from five major public hospitals (covering 57% of national acute bed capacity) for Singapore. Error bars represent the 95% confidence intervals for the mean incidence rate per 1,000,000 person-years, calculated using bootstrapping with 10,000 replicates.

Atlantic states were similarly divided into two groups by latitude. Northern Atlantic states shared a similar vibriosis etiology with the Southern Pacific region (California). A Monte Carlo Pearson χ^2^ test found no significant difference in *Vibrio* species composition between the Northern Atlantic and Southern Pacific (χ^2^ = 4.59, *p* = 0.420) (Fig. 1). In contrast to the Northern Atlantic, the Southern Atlantic states (from the Chesapeake Bay southward), exhibited a higher relative proportion of *V. vulnificus* (OR = 5.78; SE = 1.50; 95% CI = 3.23– 10.34; *z* = 6.76; *p* < 0.001) and a lower proportion of *V. parahaemolyticus* infections (OR = 0.48; SE = 0.08; 95% CI = 0.33–0.69; *z* = –4.45; *p* < 0.001) (Fig. 1). Moving south into the subtropics, Gulf Coast states were characterized by an even higher relative proportion of *V. vulnificus* infections. The odds of a *Vibrio* case being *V. vulnificus* were 1.4 times higher in the Gulf Coast than in the Southern Atlantic (OR = 1.35; SE = 0.22; 95% CI = 0.93–1.96; *z* = 1.83; *p* = 0.067). These Gulf Coast states reported some of the highest *V. vulnificus* incidence nationally, alongside Maryland and Hawaii (Fig. 2 and Supplementary Fig. 2).

Hawaii formed a distinct category and reported the highest vibriosis incidence in the United States (Fig. 2 and Supplementary Fig. 2). The incidence in Hawaii was 5.5-fold higher than the mean incidence across all other US regions (rate ratio = 5.52; SE = 0.60; 95% CI = 4.12–7.40; *z* = 15.69; *p* < 0.001), assuming the population was held constant. The state had an exceptionally high proportion of *Vibrio* infections caused by *V. alginolyticus* (53%) (Fig. 1), and the highest incidence nationwide for this species as well as for *V. vulnificus* and *V. fluvialis* (Fig. 2 and Supplementary Fig. 2). Compared to the means of other US regions, incidence in Hawaii was 15.5-fold for *V. alginolyticus* (*p* < 0.001), 16.1-fold for *V. vulnificus* cases (*p* < 0.001), and 8.2-fold for *V. fluvialis* cases (*p* < 0.001).

### Causes and incidence of vibriosis in Singapore and the United States

The species profile of *Vibrio* infections in Singapore differed markedly from that observed in the United States. Although *V. parahaemolyticus* was the most common causative agent in both countries (37% of infections), *V. fluvialis* accounted for a substantially larger proportion of infections in Singapore (27%) compared to the United States (5%). Conversely, *V. alginolyticus*, the second most common cause of vibriosis in the United States (15%), was rare in Singapore (3%). *V. furnissii*, while accounting for fewer than 0.01% of cases in the United States (42 cases from 2007–2019), in line with many other “exotic” (rare) *Vibrio* species, ranked as the fifth most common causative agent in Singapore (6%).

Overall vibriosis incidence was 2.2 times higher in Singapore than in the United States (rate ratio = 2.17; SE = 0.26; 95% CI = 1.56–3.00; *z* = 6.37; *p* < 0.001) (Fig. 3). As shown in Fig. 1, the US numbers reflected national average estimates across diverse geographic regions with substantial variation in incidence. When incidence was stratified by region, Hawaii, one of the only tropical regions in the United States along with the southern tip of Florida, reported the highest overall incidence of vibriosis. Singapore and the Northern Pacific states followed closely behind, with no statistically significant difference between them (*p* = 0.135) (Fig. 2). Singapore reported the highest incidence of *V. furnissii* and *V. fluvialis* of any region examined, and the second highest incidence of *V. cholerae*, exceeded only by the Gulf Coast state of Louisiana (*p* = 0.079) (Supplementary Fig. 2).

**Fig. 3.**
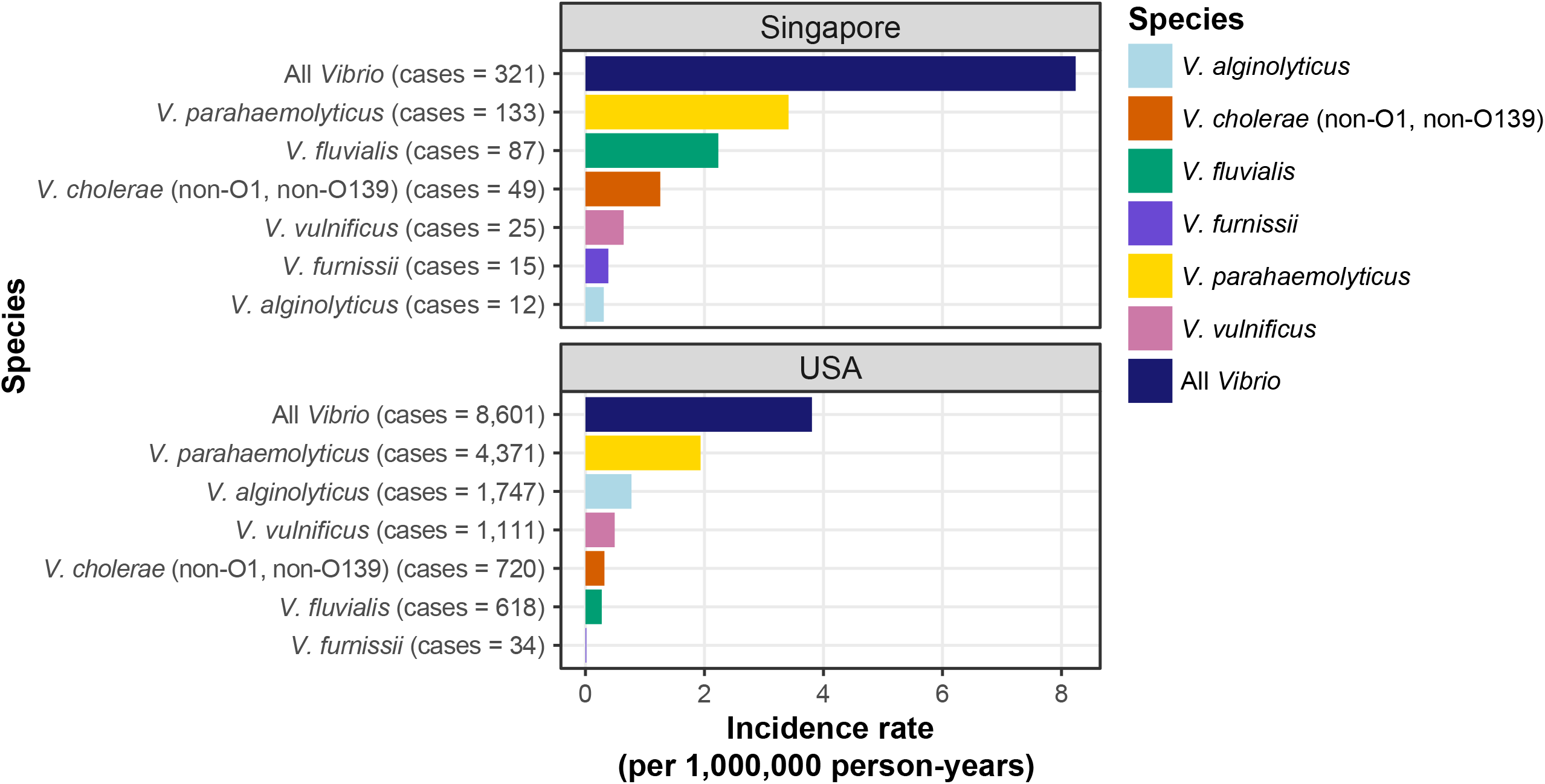
Comparison of incidence rates of vibriosis between Singapore and the United States. Incidence rates were calculated for 2013–2019 from the US COVIS database and records from five major public hospitals (covering 57% of national acute bed capacity) for Singapore.

### Temporal trends of vibriosis in Singapore and the United States

The two species that were more prevalent causes of vibriosis in Singapore than in the United States (*V. cholerae* and *V. fluvialis*) were also the fastest-rising causes of infections in the United States (Fig. 4 and Table 1). From 2007 to 2019, the incidence of *V. cholerae* increased by an average of 17% per year, a significantly higher rate than all other *Vibrio* species except *V. fluvialis* (Table 2). The latter increased by an average of 13% per year, representing the second highest rise (Table 1). Between 2007 and 2019, the number of reported infections more than tripled for *V. cholerae* and quintupled for *V. fluvialis*, whereas this number only roughly doubled for other species. In 2019, *V. cholerae* surpassed *V. vulnificus* in annual case count, marking the first time this occurred since reporting began.^7^ On the other hand, there was no statistically significant increase in vibriosis incidence in Singapore between 2013 and 2019 (*p* = 0.630) (Supplementary Fig. 3).

**Table 1.** Trends in *Vibrio* infections by species between 2007–2019 in the United States.

| Species | Ratio<br>change of<br>incidence | Lower<br>95% CI* | Upper<br>95% CI* | Z-value | Adjusted<br>p-value |
| --- | --- | --- | --- | --- | --- |
| *V. alginolyticus* | 1.090 | 1.053 | 1.128 | 4.888 | <0.001 |
| *V. cholerae* | 1.171 | 1.129 | 1.214 | 8.468 | <0.001 |
| *V. fluvialis* | 1.132 | 1.089 | 1.176 | 6.279 | <0.001 |
| *V. parahaemolyticus* | 1.088 | 1.053 | 1.126 | 4.959 | <0.001 |
| *V. vulnificus* | 1.056 | 1.020 | 1.094 | 3.055 | 0.002 |

\*CI, confidence interval

**Table 2.** Pairwise comparisons of *Vibrio* infections trends by species between 2007–2019 in the United States.

| Comparison | Ratio of<br>change<br>between<br>species | Lower<br>95%<br>CI* | Upper<br>95%<br>CI* | Z-value | Adjusted<br>p-value |
| --- | --- | --- | --- | --- | --- |
| *V. alginolyticus* / *V. cholerae* | 0.931 | 0.867 | 1.001 | −2.775 | 0.018 |
| *V. alginolyticus* / *V. fluvialis* | 0.963 | 0.894 | 1.038 | −1.414 | 0.253 |
| *V. alginolyticus* / *V. parahaemolyticus* | 1.002 | 0.935 | 1.073 | 0.062 | 0.951 |
| *V. alginolyticus* / *V. vulnificus* | 1.032 | 0.962 | 1.107 | 1.248 | 0.253 |
| *V. cholerae* / *V. fluvialis* | 1.034 | 0.959 | 1.116 | 1.247 | 0.253 |
| *V. cholerae* / *V. parahaemolyticus* | 1.075 | 1.002 | 1.154 | 2.878 | 0.018 |
| *V. cholerae* / *V. vulnificus* | 1.108 | 1.030 | 1.191 | 3.969 | <0.001 |
| *V. fluvialis* / *V. parahaemolyticus* | 1.040 | 0.966 | 1.119 | 1.492 | 0.253 |
| *V. fluvialis* / *V. vulnificus* | 1.071 | 0.994 | 1.154 | 2.582 | 0.025 |
| *V. parahaemolyticus* / *V. vulnificus* | 1.030 | 0.961 | 1.105 | 1.207 | 0.253 |

\*CI, confidence interval

**Fig. 4.**
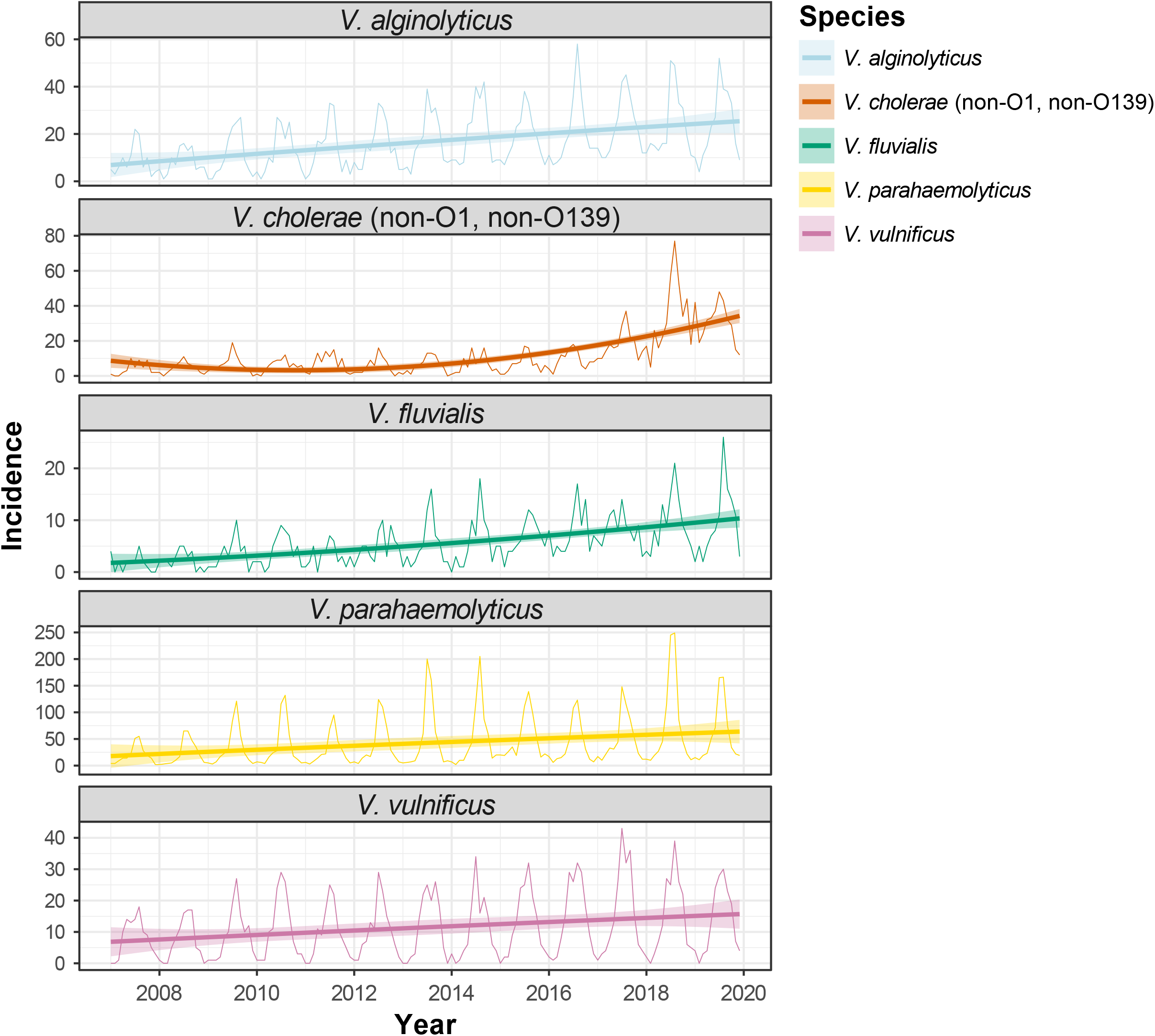
Cases of vibriosis by causative species in the United States, 2007–2019. Monthly cases of vibriosis were calculated to illustrate seasonality. A smoothed quadratic regression line (second degree polynomial) was fitted for each species, with shaded areas representing the 95% confidence intervals.

### Seasonal patterns for vibriosis in the mainland United States

Most infections peaked during the summer months in the United States (June to August) (Fig. 5A) with Hawaii displaying a similar seasonal pattern, albeit less pronounced (Fig. 5B). However, seasonal patterns were not uniform across species in the United States.

**Fig. 5.**
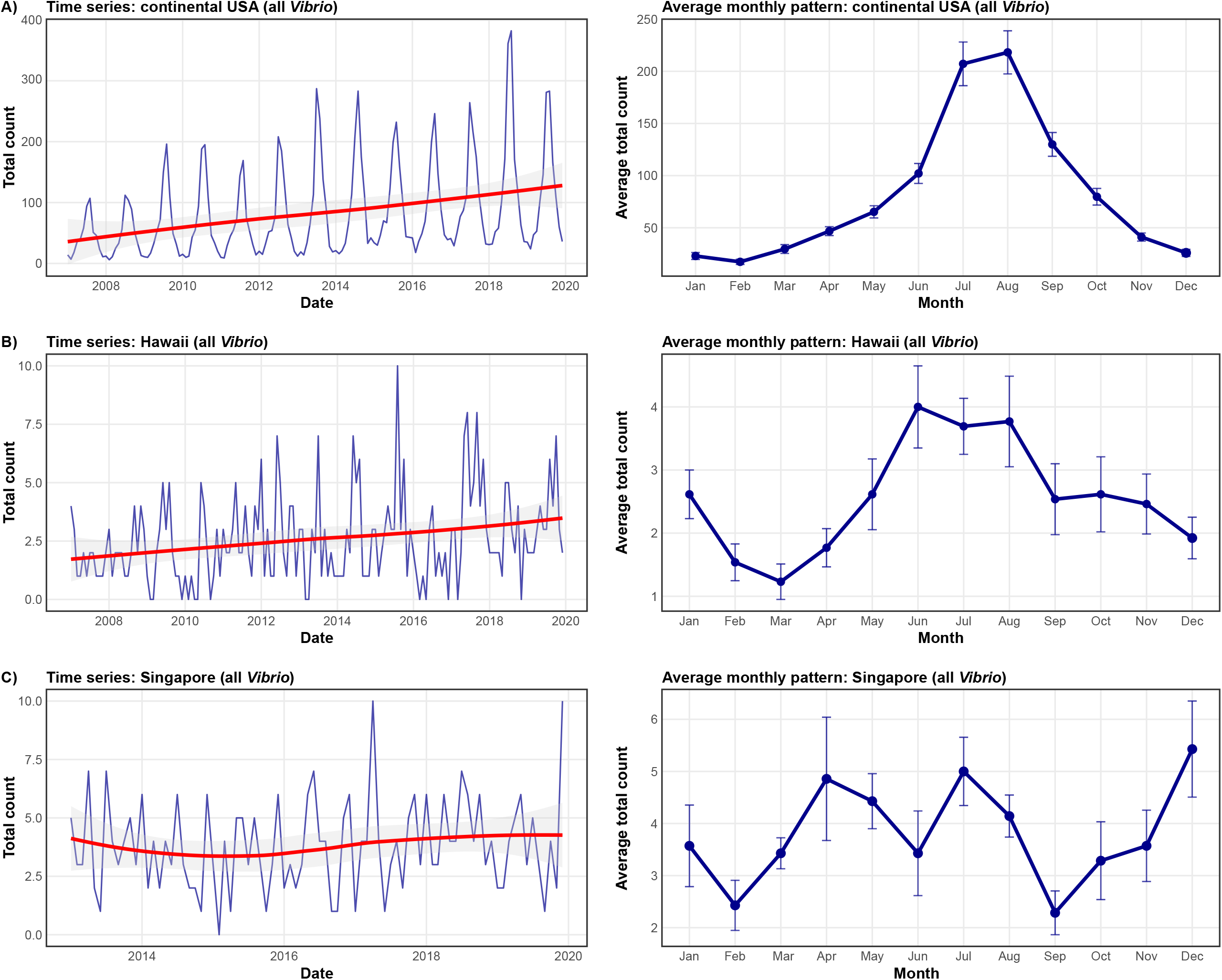
Change in reported vibriosis cases across regions with different climates. Yearly and monthly patterns are shown for the continental United States (temperate– subtropical) (A), Hawaii (tropical) (B), and Singapore (equatorial) (C). Smoothed mean lines (left panels) were calculated using LOESS regression (span = 1.0), with shaded areas representing the 95% confidence intervals. Error bars (right panels) represent the standard error of the mean monthly case counts across years.

The percentage of US cases occurring during the summer was 40.9% for *V. fluvialis*, 41.7% for *V. cholerae*, 44.3% for *V. alginolyticus*, 52.3% for *V. vulnificus*, and 60.2% for *V. parahaemolyticus*. These values represented an increase in cases during the summer relative to a comparable period during the rest of the year, by a factor of 2.1 for *V. fluvialis*, 2.2 for *V. cholerae*, 2.4 for *V. alginolyticus*, 3.3 for *V. vulnificus*, and 4.5 for *V. parahaemolyticus*, underscoring the variation in seasonal amplification across *Vibrio* species. The more pronounced seasonality of *V. vulnificus* and *V. parahaemolyticus* infections was also reflected in their seasonal ratios (peak/trough) (Supplementary Fig. 4).

Regional differences in seasonality were also evident (Fig. 6). The Gulf Coast accounted for most cases of vibriosis in spring (March to May), when the water temperatures there are relatively higher than in other regions.^8^ Hawaii contributed a greater proportion of cases during winter (December to February), consistent with its relatively stable sea surface temperatures year-round.^8^ In contrast, the northern regions (both Pacific and Atlantic coasts) contributed most prominently in the summer. Non-coastal regions maintained a relatively constant proportion of the national burden throughout the year.

**Fig. 6.**
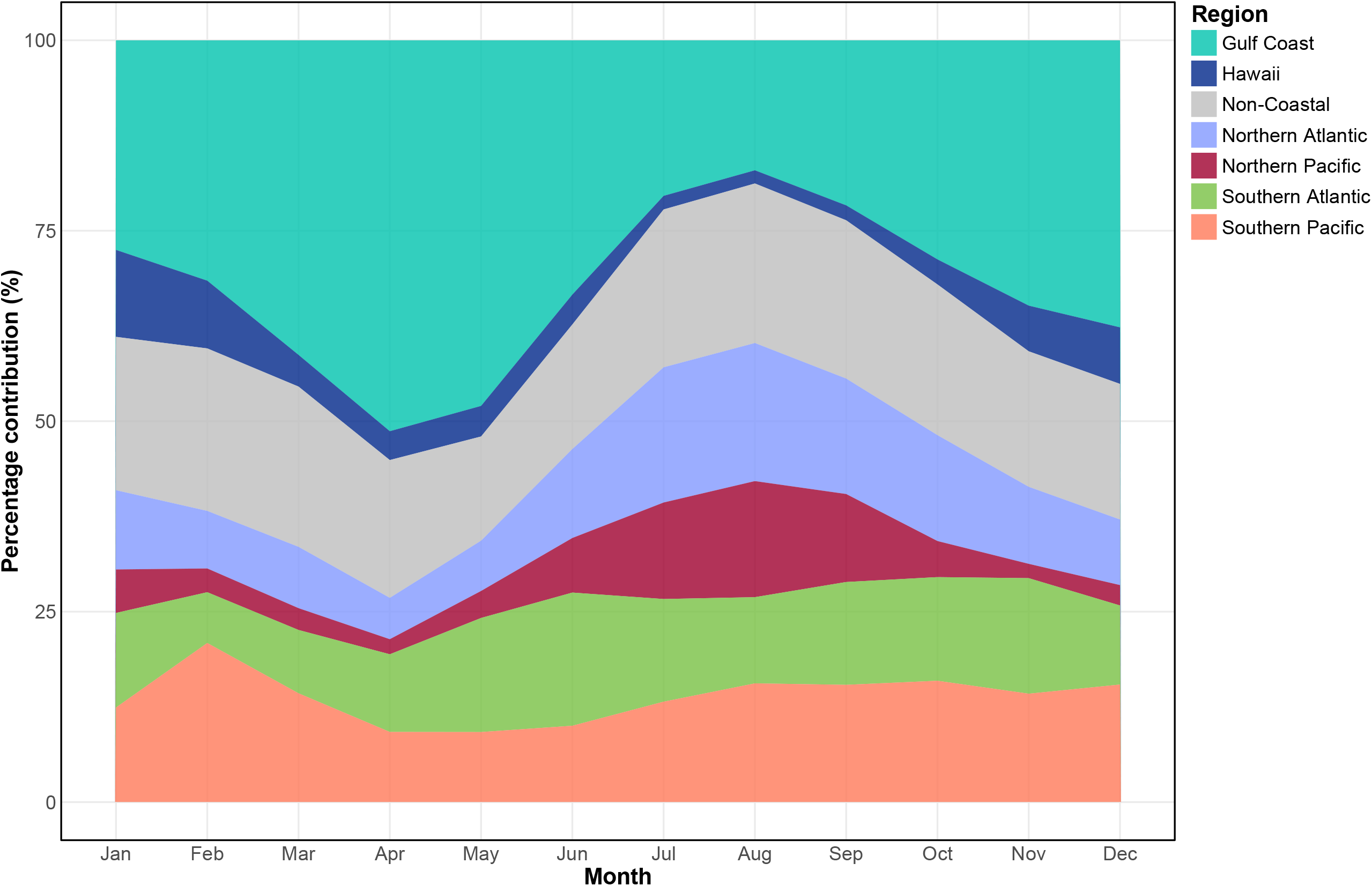
Regional contributions to vibriosis in the United States. Monthly proportions of reported vibriosis cases (all species combined) from 2007–2019 are shown for seven US regions (defined in Fig. 1B). Stacked area plots illustrate the dynamic contribution of each region to the national seasonal pattern, with areas summing to 100% each month.

### Vibriosis incidence trends and seasonality patterns in three unique climates

The mainland United States has a largely temperate climate, experiencing a wide range of coastal water temperatures over a year. Differences between summer and winter monthly mean water temperatures typically differ by around 20°C in any given region (and at least 10°C for regions with the narrowest range such as Alaska or Southern Florida).^8^ There, vibriosis incidence is increasing rapidly, with a pronounced seasonality (Fig. 5A). Hawaii, with a tropical climate and relatively stable average water temperatures (23–28°C),^8^ experiences modest increases in vibriosis incidence in the summer and a consistent increase in cases throughout the last decade, although not to the same extent as the mainland (Fig. 5B). Singapore, with an equatorial monsoonal climate, displays consistent but small seasonal variation in sea surface temperature (28–31°C).^9^ There, vibriosis does not follow the temperate pattern of increased infections in summer, and no significant increase in infections can be seen between 2013 and 2019 (Fig. 5C). However, when looking at the cumulative results of culture-independent diagnostic tests (CIDTs) for Singapore hospitals from the start of their use in 2017 until 2024, the monsoonal seasonality of vibrio infections is clear (Supplementary Fig. 5). The larger amount of data available for CIDTs and increase in signal by using cumulative counts (as opposed to culture-confirmed cases monthly averages used in this study) shows the clear relation between water temperature and infections, even at the elevated levels seen at the equator.

## DISCUSSION

The fact that equatorial Singapore showed no statistically significant increase in vibriosis despite higher incidence than the United States might be linked to the negligible water temperature increase in the last 50 years (0.00576°C per year, <0.3°C) (Supplementary Fig. 6). As a point of comparison, average water temperature in the North Atlantic has increased five times faster during this time period (∼1.5°C).^10^ Despite the lack of overall increase in cases in Singapore, the seasonal variation of water temperature resulting from the Northeast and Southwest monsoons is clearly correlated with the number of cases when using cumulative case data available, which peak with the maximum monthly mean water temperature of 30.5°C (Supplementary Fig. 5). This suggests that the relationship between water temperature and incidence persists even beyond 30°C, and it remains unclear whether this trend will plateau at higher temperatures.

While water temperature is a major driver, there are other factors known to affect the incidence of vibriosis. These include population density along the coast, types and amounts of seafood consumed, frequency and nature of recreational water activities, and the demographic and health characteristics of exposed populations.^3^ The propensity for extreme weather events can also drive risk in some areas like the coast of the Baltic Sea (heatwaves) and the Gulf Coast of the United States (hurricanes).^11,12^ States like Texas and Florida, for example, have extensive coastlines and large populations living along them,^13^ which may contribute to their consistently high infection counts. Host susceptibility also plays an important role. Adult males with underlying health conditions, particularly liver disease, are at increased risk for severe *V. vulnificus* infections, which are associated with disproportionately high mortality.^14^ The salinity of coastal waters, linked to rainfall patterns and freshwater input from rivers, is known to affect *Vibrio* species differentially.^11^ Additionally, biotic environmental contributors such as the frequency and type of zooplankton and phytoplankton blooms, or the concentration of dissolved and suspended organic carbon in water, may also influence species-specific patterns of infection, although these factors have yet to be directly investigated.^3^

The available data suggests that different *Vibrio* species are differentially influenced by this complex combination of environmental, ecological, and social factors. For instance, the high relative proportion of infections caused by *V. fluvialis* in Singapore, compared with the United States, suggests that equatorial climates may favor its persistence or transmission. The elevated incidence of *V. fluvialis* in warm-water US states such as Louisiana and Hawaii supports this hypothesis. Meanwhile, *V. cholerae* infections, whose incidence is rising more rapidly than any other species, make up a larger proportion of cases in non-coastal US states compared to coastal states. This may be due to the ability of the species to persist in brackish and even freshwater environments, as shown in European studies (Lake Neusiedl, Austria)^15^ and limited US reports of exposures in rivers and lakes in Michigan, Missouri, Texas, and Arizona,^16^ combined with the fact that *V. cholerae* is predominantly foodborne (Supplementary Fig. 7) and inland states rely on seafood distributed through national supply chains originating from coastal ports.^17^ Its incidence is also highest in Louisiana and Singapore, both of which have warm waters.^8,9^ Additional factors, such as seafood import networks, host susceptibility, and surveillance capacity, may shape observed patterns.^3^

Exposure may also explain why waters cooler than that of Singapore support higher incidences of certain species. For example, *V. alginolyticus* and *V. vulnificus* have higher incidence in Hawaii than in all other regions. There, sea surface temperatures are lower (23– 28°C) than in Singapore (28–31°C).^8,9^ Because these species are commonly associated with wound and soft tissue infections linked to seawater exposure (Supplementary Fig. 7), differences in recreational water activity may contribute. While at least 80% of Americans aged 15 and older report being able to swim without assistance, just over 60% of Singaporeans do, suggesting potentially lower water exposure in the latter.^18^ However, environmental, social, or behavioral factors, such as raw oyster consumption, which is far more common in the United States, likely contribute as well.^3^

As Singapore is a nation-state with a relatively modest population, we also examined available data from neighboring Malaysia to assess for similar patterns. Vibriosis trends in Singapore were indeed consistent with limited findings from Malaysia, which shares similar coastal waters and is among Singapore’s major seafood suppliers.^19,20^ As in Singapore, the number of cases remained relatively stable year to year, with *V. parahaemolyticus* and *V. fluvialis* identified as the predominant causative agents.^19^ This regional coherence supports the robustness of the Singapore findings and suggests that similar environmental and ecological conditions shape *Vibrio* species distribution in the region.

To our knowledge, this study includes one of the most comprehensive national datasets available outside the United States. However, one important limitation is that only 57% of total acute bed capacity within Singapore was captured.^21^ Incidence estimates for Singapore therefore likely underrepresent the true national burden. Adjusting for this sampling coverage and assuming similar case detection rates across all facilities, if all national data had been available, true incidence could be nearly double the current reported values. In such a scenario, the overall vibriosis incidence of Singapore would be comparable to Hawaii and exceed all US states (including Hawaii) for *V. furnissii, V. cholerae*, and *V. fluvialis*. We also chose to limit the data to the end of 2019, as vibriosis incidence significantly dropped in the years of the COVID-19 pandemic, likely due to both less exposure but also increased underreporting and underdiagnosis, which would have biased the results.

Overall, these findings highlight the strong climate dependence of vibriosis, suggesting that incidence in the United States is likely to continue rising at a fast pace and that infection patterns are tropicalizing, becoming more similar to those observed in tropical or equatorial climates such as Singapore. Even in the warm waters of this equatorial island-nation, infection numbers remain strongly associated with water temperature, with no indications of plateauing even above 30°C. The limit in this temperature-infection relationship is currently unknown, but given that optimal growth temperature of most of the pathogenic *Vibrio* discussed here is approximately 37°C, it is not likely that increase in infection will slow down in the near future. The apparent plateau observed in Singapore is likely due to a slower warming rate (about five times slower than the colder waters of the Northern Atlantic), not to a breakdown of the temperature-infection positive correlation at high temperatures.

## METHODS

### Data acquisition

This retrospective study analyzes the epidemiology of vibriosis over a minimum 10-year period, corresponding to the duration of available surveillance data in Singapore. The dataset comprised 9,324 aggregated monthly observations, encompassing case counts from different geographical regions of the United States and Singapore. Each observation included the residence region (state), *Vibrio* species designation, temporal variables (year and month), total case counts, population denominators, and pre-calculated incidences.

The United States is one of the few countries where cholera and vibriosis are nationally notifiable diseases. The CDC maintains the COVIS database, used for reporting *Vibrio* cases in the United States since 1988.^7^ Public health officials interview patients using a standardized case report form and submit findings to the CDC, including patient demographics, risk factors, and bacterial information. The period of data collection and analysis extends from the beginning of 2007, when *Vibrio* cases first became nationally notifiable, through 2019, due to potential disruptions in case reporting and processing during the COVID-19 pandemic.

In Singapore, as only limited surveillance of vibriosis is done at a national level (cholera is the only notifiable *Vibrio*-related disease),^22^ data were retrieved from five participating public acute hospital laboratories: the Changi General Hospital (CGH), Singapore General Hospital (SGH), National University Hospital (NUH), Ng Teng Fong General Hospital (NTFGH), and Tan Tock Seng Hospital (TTSH). These five major public hospitals account for approximately 65% of acute public hospital beds and 57% of total acute bed capacity in Singapore.^21^ As with US laboratories, there is some variability in pathogen identification methods among these institutions. Singapore data were collected from the beginning of 2013 through 2023. For comparisons with the United States, only data from 2013 to 2019 were used to ensure matching timeframes. All records were de-identified. The use of Singapore hospital data was approved by the National Health Group Domain Specific Review Board on February 28, 2024 (reference number 2023/00917), with waiver of patient consent obtained. The same case definition was used for vibriosis in the USA and Singapore and is described in supplementary materials.

### Data variables

The top five causative agents (species) for vibriosis in each country were identified. This resulted in six species being included in the analysis, as *V. furnissii* ranked in the top five in Singapore but not in the United States, and *V. alginolyticus* was among the top five in the United States but not in Singapore. *V. cholerae* O1/O139 were excluded from the analysis, as these are likely to represent imported cases of pandemic cholera.^7^ In this study, *V. cholerae* refers exclusively to non-O1/non-O139 (non-pandemic) strains, which were likely acquired locally. The six species included in the final analysis were *V. alginolyticus, V. cholerae* (non-O1/non-O139), *V. fluvialis, V. furnissii, V. parahaemolyticus*, and *V. vulnificus*. In addition to species-specific counts, an aggregate ‘All *Vibrio*’ category was created to represent total infections, with population-normalized rates (cases per 1,000,000 person-years) to enable comparison across regions differing in population size.

To account for inconsistencies in date recording across databases and potential delays between specimen collection and reporting, only the month and year were extracted for analysis. To facilitate comparison between the United States (COVIS) and Singapore, samples were classified into three categories based on likely origin: foodborne/likely foodborne, non-foodborne/likely non-foodborne, and unknown. Seasons were based on the Northern Hemisphere calendar, where both countries are located (Autumn: September to November; Winter: December to February; Spring: March to May; Summer: June to August).

### Regional differences in species proportions

Infection counts were individually transformed into proportions for each US state and for Singapore to determine the share of total *Vibrio* infections caused by each species. To assess geographic differences in species profiles, Bray–Curtis dissimilarity was calculated on these proportions, and states were grouped using average-linkage hierarchical clustering.^23^ Based on the resulting clusters, seven US geographical regions were defined: Northern Atlantic, Southern Atlantic, Northern Pacific, Southern Pacific, Gulf Coast, Hawaii, and Non-Coastal.

To test for differences in species proportions across regions, daily counts for each *Vibrio* species were paired with total daily *Vibrio* counts in each region and converted to species-specific proportions. These were modeled using a weighted quasi-binomial logistic regression (weights = monthly totals), with region as the sole predictor. Fitted contrasts provided odds ratios quantifying how much more (or less) likely infection with each species was in one region compared to another. Overall species composition was assessed by pooling species-level case counts within each region and expressing them as a percentage of that region’s total *Vibrio* burden. Differences in species composition between regions were tested using a Pearson χ^2^ test with *p*-values obtained via 1,000,000 Monte Carlo replicates.^24^ An alpha level of 0.05 was used for statistical significance.

### Incidence estimation and temporal trends

Crude incidences were calculated using population estimates from the respective national censuses.^25,26^ To determine annual incidence by species, state, and region, annual case counts from 2013 to 2019 were summed and divided by the corresponding population, yielding incidence per 1,000,000 person-years. Sampling uncertainty was quantified using a non-parametric bootstrap (10,000 replicates) within each species–region/state stratum.^27^

A negative binomial model which incorporates a gamma-distributed random effect to account for overdispersion was used, allowing the variance to exceed the mean and better captures variability in incidence.^28^ An offset term was included to adjust for differing population sizes, ensuring the model estimates true incidences rather than raw counts. To reduce the risk of Type I errors (false positives) due to multiple comparisons, *p*-values were adjusted using the Benjamini–Hochberg (BH) procedure to control the false discovery rate (FDR), the expected proportion of false positives among rejected hypotheses.^29^ The annual ratio of change was estimated for each species using the following model structure: *Incidence ∼ Species × Time since first incidence + offset(log(Population)), family = Negative Binomial(link = log)*.

### Seasonality analysis

To examine species-specific changes in incidence across the United States from 2007 to 2019, monthly incidence values were calculated to illustrate seasonality. A smoothed quadratic regression line (second-degree polynomial) was fitted for each species, with 95% confidence intervals. For each defined US region, unadjusted monthly values were used to generate smoothed means using Locally Estimated Scatterplot Smoothing (LOESS) regression (span = 1.0),^30^ also with 95% confidence intervals. The same approach was applied to the Singapore data from 2013–2019, the period for which data overlapping with the US data timeframe was available.

To assess the influence of climate variability, temporal trends were compared across three climatically distinct regions: continental United States (including Alaska; temperate– subtropical), Hawaii (tropical), and Singapore (equatorial). LOESS smoothing (span = 1.0) was used to capture both long-term and seasonal variation.^30^ For within-year characterization, average monthly case counts were calculated with associated standard errors, determined as the standard deviation divided by the square root of the number of observations per month. These analyses were based on total *Vibrio* cases to capture the overall disease burden.

### Regional contribution analysis

To analyze geographic contributions to the national seasonal signal in the United States, the proportional contribution of each state to total monthly case counts was calculated as *(regional cases / total US cases) × 100*. Regions were ranked by their average annual contribution, and results were visualized using stacked area plots. These plots displayed the dynamic monthly contribution of regions to the overall US seasonal pattern, with regions stacked to sum to 100% for each month.

## DATA AVAILABILITY

Restrictions apply to the availability of these data. Data were obtained under data sharing agreements from contributing surveillance sites and can only be shared by contributing organizations with their permission.

## ACKNOWLEDGEMENTS

We would like to thank the Biotechnology and Biological Sciences Research Council BB/Y514068/1 (International Institutional Awards Quadram) for supporting a workshop to discuss this work, as well as the National University of Singapore Saw Swee Hock School of Public Health for covering publication fees. Seawater temperature measurements in Singapore were obtained from the Marine Environmental Sensing Network (https://ombak.mesn.sg).

## AUTHOR CONTRIBUTIONS

CCN: methodology, formal analysis, investigation, data curation, visualization, writing – original draft. EDH: methodology, formal analysis, data curation, visualization. FDO: data curation, visualization, writing – review & editing. CBA: conceptualization, writing – review & editing. MHa: conceptualization, writing – review & editing. MHu: resources, writing – review & editing. BMHK: data curation, visualization, writing – review & editing. PM: data curation, resources. TYE: resources. CSLW: resources. CKL: resources. SGDC: resources. CY: resources. TB: resources. SM: methodology, formal analysis, data curation, visualization. JMU: conceptualization, writing – review & editing. AEM: conceptualization, writing – review & editing. CL: resources, conceptualization, writing – review & editing. YFB: conceptualization, methodology, visualization, resources, data curation, writing – original draft, writing – review & editing, supervision, project administration. CCN, EDH, SM, and YFB accessed and verified the underlying data reported in the manuscript. All authors confirm that they had full access to all the data in the study and accept responsibility to submit for publication.

## REFERENCES

1. Baker-Austin, C., Trinanes, J., Gonzalez-Escalona, N. & Martinez-Urtaza, J. Non-cholera vibrios: the microbial barometer of climate change. Trends Microbiol 25, 76–84, doi:10.1016/j.tim.2016.09.008 (2017).

2. Islam, M. T., Alam, M. & Boucher, Y. Emergence, ecology and dispersal of the pandemic generating Vibrio cholerae lineage. Int Microbiol 20, 106–115, doi:10.2436/20.1501.01.291 (2017).

3. Baker-Austin, C. et al. Vibrio spp. infections. Nat Rev Dis Primers 4, 1–19, doi:10.1038/s41572-018-0005-8 (2018).

4. Johnson, C. N. Influence of environmental factors on Vibrio spp. in coastal ecosystems. Microbiol Spectr 3, VE-0008-2014, doi:10.1128/microbiolspec.VE-0008-2014 (2015).

5. Amato, E. et al. Epidemiological and microbiological investigation of a large increase in vibriosis, northern Europe, 2018. Euro Surveill 27, 2101088, doi:10.2807/1560-7917.ES.2022.27.28.2101088 (2022).

6. Newton, A., Kendall, M., Vugia, D. J., Henao, O. L. & Mahon, B. E. Increasing rates of vibriosis in the United States, 1996–2010: review of surveillance data from 2 systems. Clin Infect Dis 54, S391–S395, doi:10.1093/cid/cis243 (2012).

7. US Centers for Disease Control and Prevention. Cholera and other Vibrio illness surveillance system, <https://www.cdc.gov/vibrio/php/surveillance/index.html> (May 14, 2024).

8. National Centers for Environmental Information. Coastal water temperature guide, <https://www.ncei.noaa.gov/access/coastal-water-temperature-guide/all_table.html> (2025).

9. Mubarak, M., Rifardi, R., Nurhuda, A., Syaputra, R. F. & Retnawaty, S. F. Sea surface temperature (SST) and rainfall trends in the Singapore Strait from 2002 to 2019. Indones J Geogr 54, 55–61, doi:10.22146/ijg.68738 (2022).

10. Vezzulli, L. et al. Climate influence on Vibrio and associated human diseases during the past half-century in the coastal North Atlantic. Proc Natl Acad Sci USA 113, E5062–E5071, doi:10.1073/pnas.1609157113 (2016).

11. Baker-Austin, C. et al. Stemming the rising tide of Vibrio disease. Lancet Planet Health 8, e515–e520, doi:10.1016/S2542-5196(24)00124-4 (2024).

12. Brumfield, K. D. et al. Genomic diversity of Vibrio spp. and metagenomic analysis of pathogens in Florida Gulf coastal waters following Hurricane Ian. mBio 14, e0147623, doi:10.1128/mbio.01476-23 (2023).

13. National Oceanic and Atmospheric Administration. National coastal population report: population trends from 1970 to 2020, <https://oceanservice.noaa.gov/facts/coastal-population-report.pdf> (March 19, 2013).

14. Hast, M. et al. Vibrio vulnificus epidemiology and risk factors for mortality in the United States, 2000-2022. Infect Dis (Lond), 1–12, doi:10.1080/23744235.2025.2559883 (2025).

15. Pretzer, C. et al. High genetic diversity of Vibrio cholerae in the European lake Neusiedler See is associated with intensive recombination in the reed habitat and the long-distance transfer of strains. Environ Microbiol 19, 328–344, doi:10.1111/1462-2920.13612 (2017).

16. Crowe, S. J. et al. Vibriosis, not cholera: toxigenic Vibrio cholerae non-O1, non-O139 infections in the United States, 1984–2014. Epidemiol Infect 144, 3335–3341, doi:10.1017/S0950268816001783 (2016).

17. Ferreira, J. P., Garlock, T., Court, C. D., Anderson, J. L. & Asche, F. The economic contribution of U.S. seafood imports throughout the value chain: a sectorial and species-specific analysis. Mar Policy 169, 106375, doi:10.1016/j.marpol.2024.106375 (2024).

18. Borgonovi, F., Seitz, H. & Vogel, I. Swimming skills around the world: evidence on inequalities in life skills across and within countries, <https://www.oecd.org/content/dam/oecd/en/publications/reports/2022/11/swimming-skills-around-the-world_ca0372da/0c2c8862-en.pdf> (September 26, 2023).

19. Hassan, M. et al. Distribution, prevalence, and antibiotic susceptibility profiles of infectious noncholera Vibrio species in Malaysia. J Trop Med 2023, 2716789, doi:10.1155/2023/2716789 (2023).

20. Singapore Food Agency. Singapore food statistics 2024, <https://www.sfa.gov.sg/docs/default-source/publication/sg-food-statistics/singapore-food-statistics-2024.pdf> (June 5, 2025).

21. Singapore Ministry of Health. Beds in inpatient facilities and places in non-residential long-term care facilities, <https://www.moh.gov.sg/others/resources-and-statistics/beds-in-inpatient-facilities-and-places-in-non-residential-long-term-care-facilities> May 26, 2025).

22. Communicable Diseases Agency. Infectious Diseases Act, <https://www.cda.gov.sg/public/infectious-diseases-act> (April 11, 2025).

23. Clarke, K. R., Somerfield, P. J. & Gorley, R. N. Clustering in non-parametric multivariate analyses. J Exp Mar Biol Ecol 483, 147–155, doi:10.1016/j.jembe.2016.07.010 (2016).

24. Waller, L. A., Smith, D., Childs, J. E. & Real, L. A. Monte Carlo assessments of goodness-of-fit for ecological simulation models. Ecol Modell 164, 49–63, doi:10.1016/S0304-3800(03)00011-5 (2003).

25. Singapore Department of Statistics. Population dashboard, <https://www.singstat.gov.sg/find-data/search-by-theme/population/population-and-population-structure/visualising-data/population-dashboard> (February 3, 2025).

26. United States Census Bureau. National population totals and components of change: 2020–2024, <https://www.census.gov/data/tables/time-series/demo/popest/2020s-national-total.html> (December, 2024).

27. Briggs, A. H., Wonderling, D. E. & Mooney, C. Z. Pulling cost-effectiveness analysis up by its bootstraps: a non-parametric approach to confidence interval estimation. Health Econ 6, 327–340, doi:10.1002/(sici)1099-1050(199707)6:4<327::aid-hec282>3.0.co;2-w (1997).

28. Linden, A. & Mäntyniemi, S. Using the negative binomial distribution to model overdispersion in ecological count data. Ecology 92, 1414–1421, doi:10.1890/10-1831.1 (2011).

29. Benjamini, Y. & Hochberg, Y. Controlling the false discovery rate: a practical and powerful approach to multiple testing. J R Stat Soc Series B Stat Methodol 57, 289–300, doi:10.1111/j.2517-6161.1995.tb02031.x (2018).

30. Cleveland, W. S. & Devlin, S. J. Locally weighted regression: an approach to regression analysis by local fitting. J Am Stat Assoc 83, 596–610, doi:10.1080/01621459.1988.10478639 (1988).

