## Supplementary Information for "Lessons from the equator: a window on the future of vibriosis in a warming Earth"

**This supplementary file includes:**

- Supplementary Figures 1–7
- Supplementary Methods
- References

### SUPPLEMENTARY FIGURES

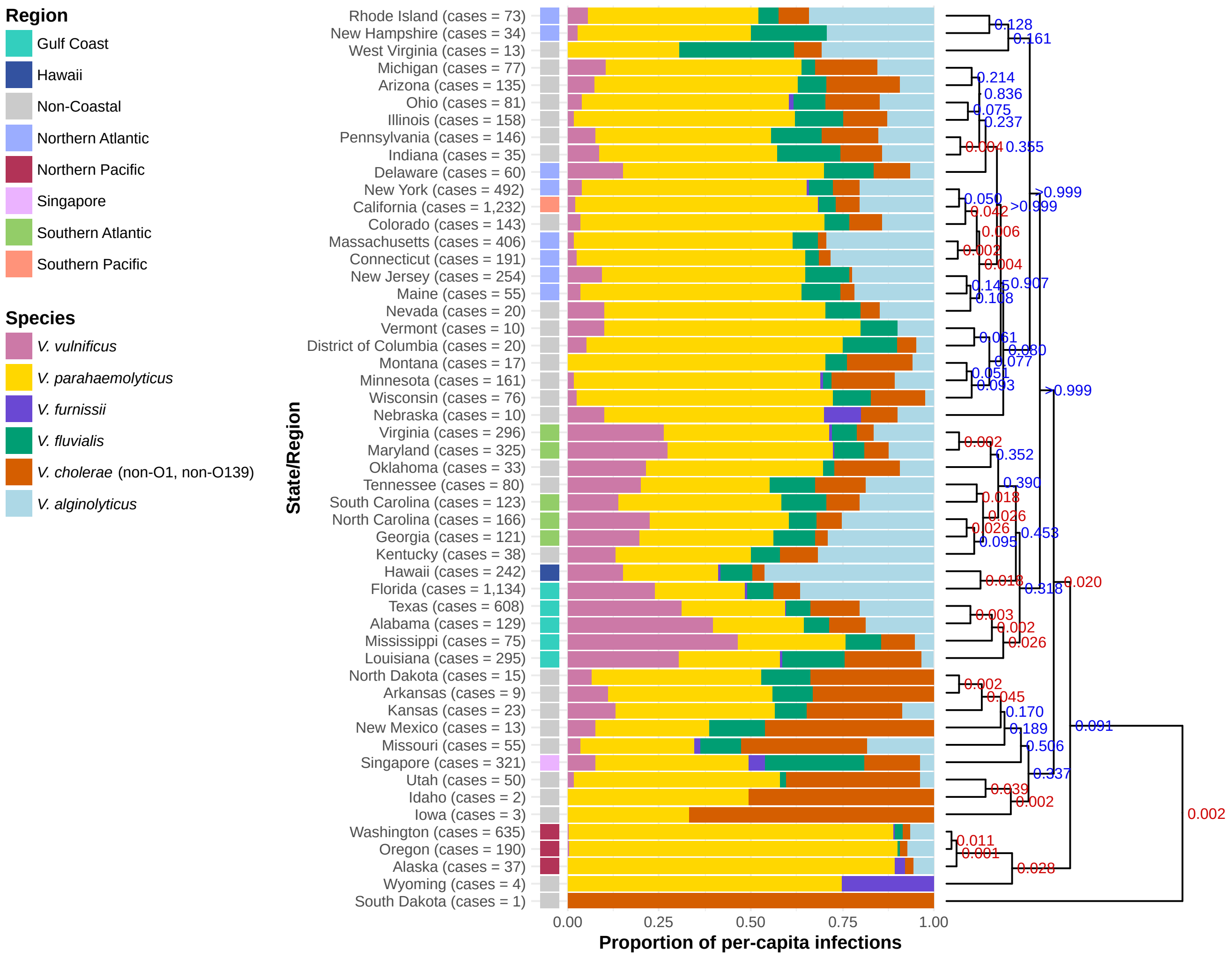


**Supplementary Fig. 1. Relative proportion of Vibrio infections by causative species across US states and Singapore.** US states and Singapore were grouped with agglomerative hierarchical clustering based on Bray–Curtis dissimilarities of their species-level relative-abundance profiles, using the average-linkage (UPGMA) method to merge clusters. The *p*-values test the null hypothesis that the cluster does not exist, with red ($p \leq 0.05$) denoting statistically significant clusters and blue ($p > 0.05$) indicating non-significant groupings.

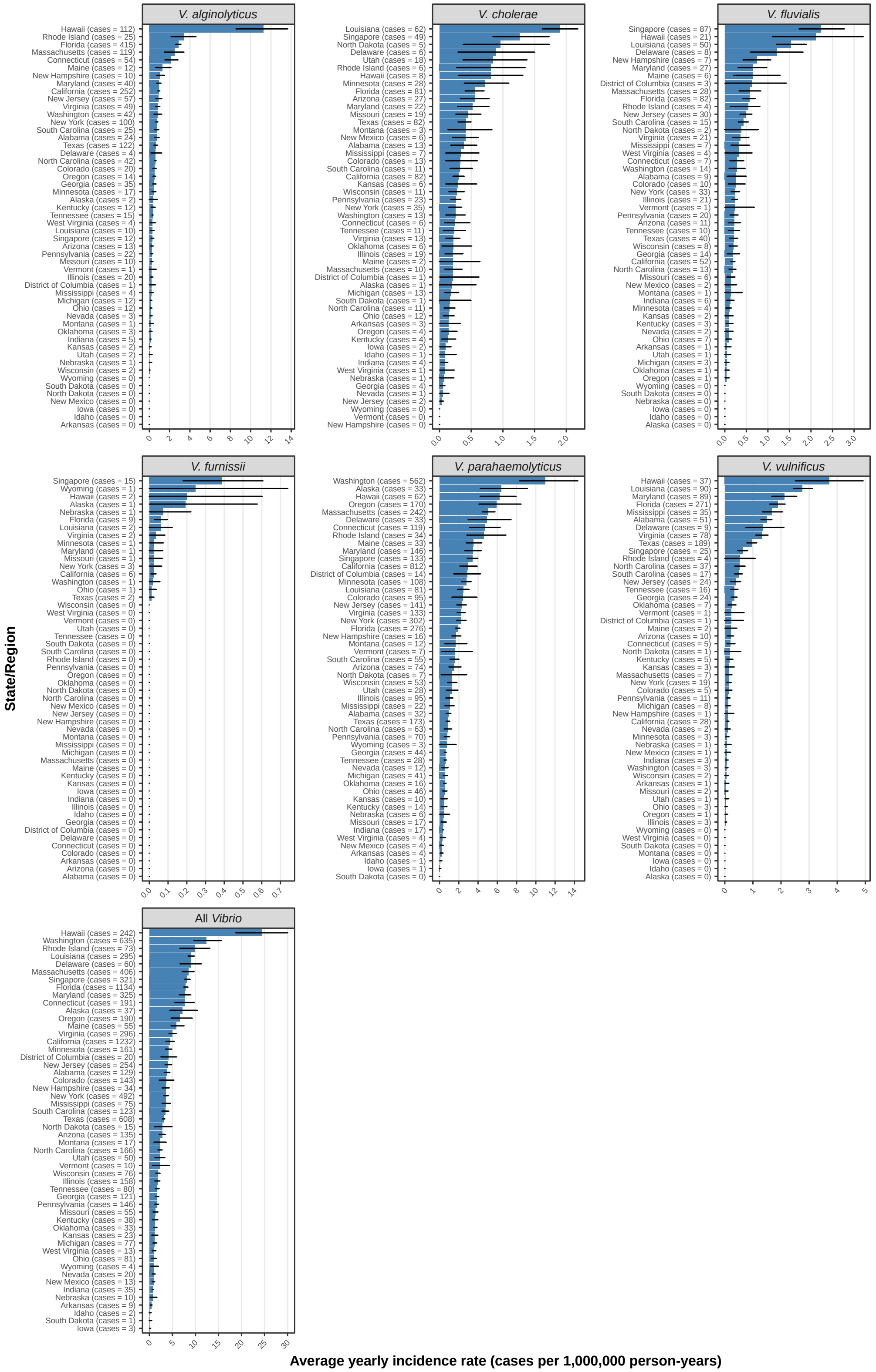


**Supplementary Fig. 2. Incidence rates of vibriosis by causative species across US states and Singapore.** Incidence rates were calculated for 2013–2019 from the US Cholera and Other Vibrio Illness Surveillance (COVIS) database and records from five major public hospitals (covering 57% of national bed capacity) for Singapore. Error bars represent the 95% confidence intervals for the mean incidence rate per 1,000,000 person-years, calculated using bootstrapping with 10,000 replicates.

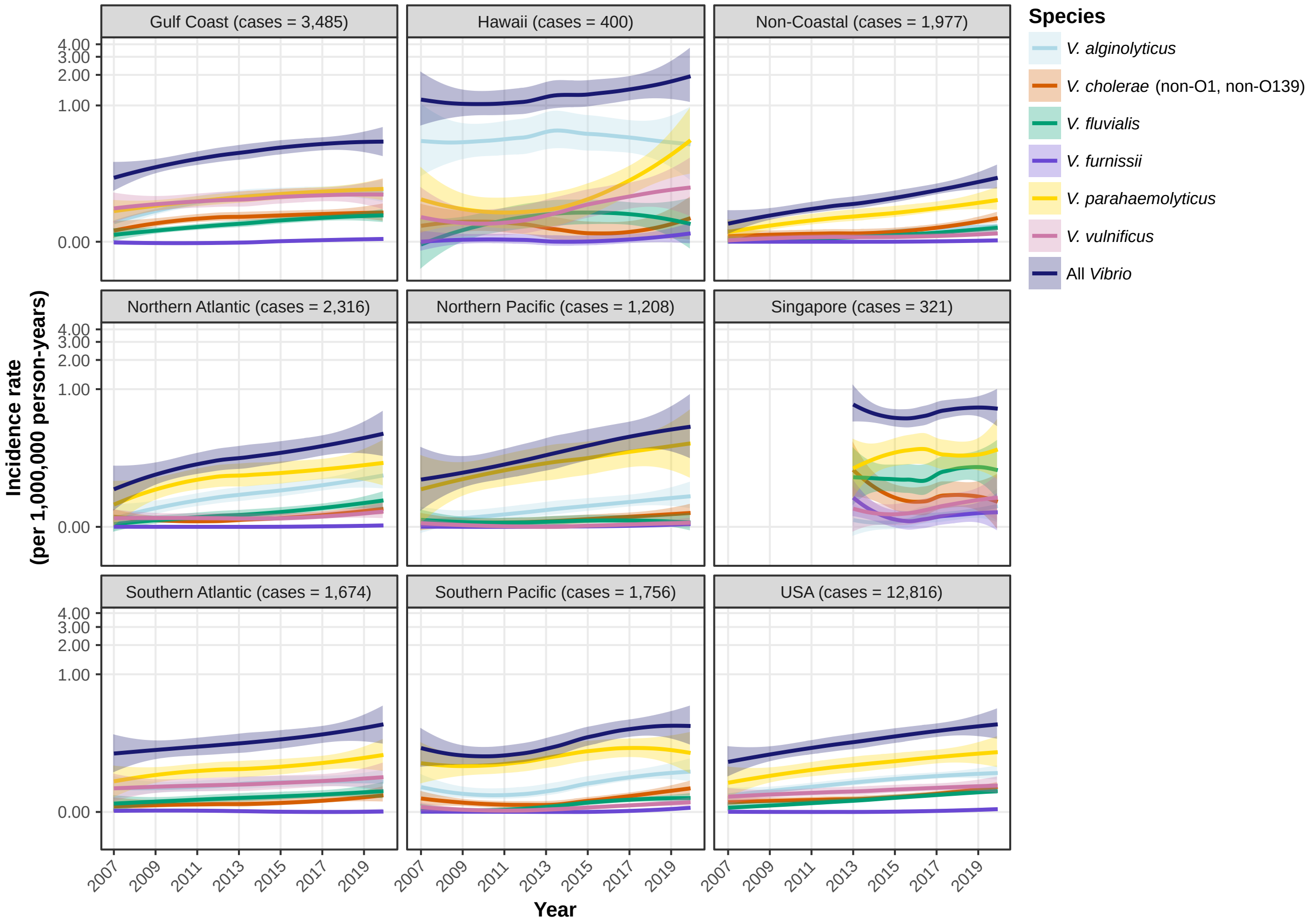


**Supplementary Fig. 3. Incidence rates of vibriosis in different US regions and Singapore.** Smoothed mean lines were calculated using LOESS regression (span = 1.0), with shaded areas showing the 95% confidence intervals.

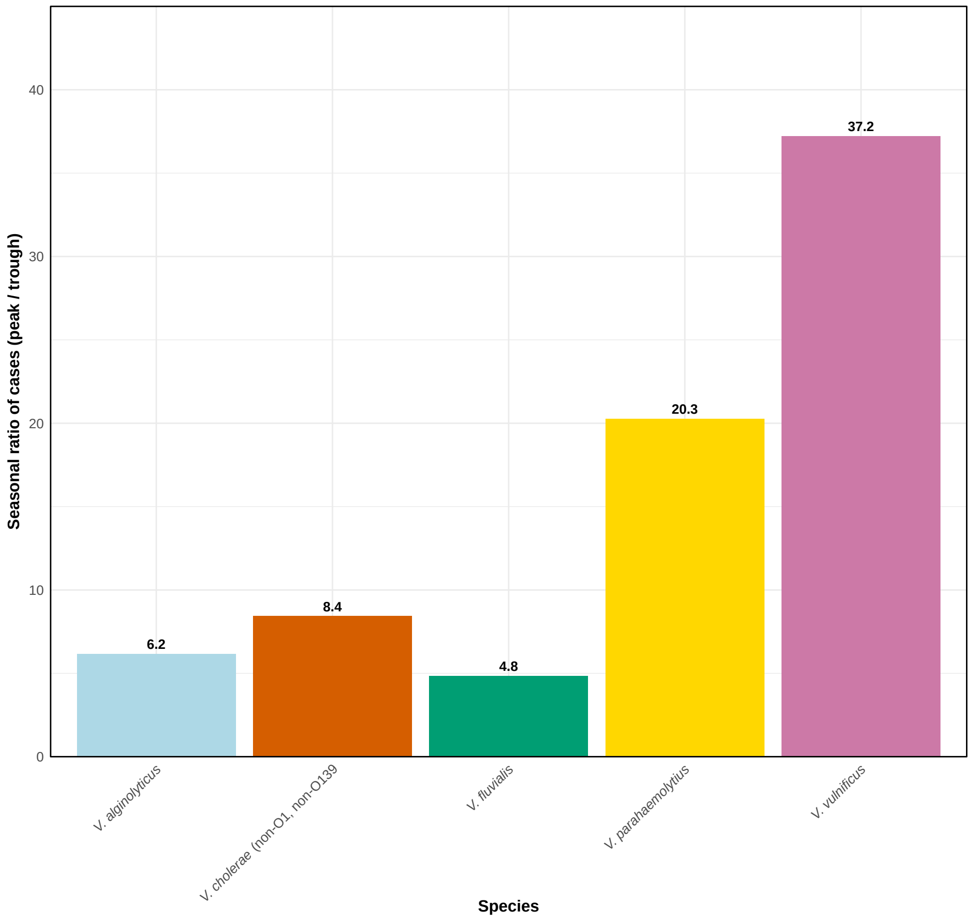


**Supplementary Fig. 4. Seasonal ratio of Vibrio infections between high and low seasons.** Ratios were calculated by comparing mean monthly incidence during the high season (summer/peak) with that during the low season (winter/trough).

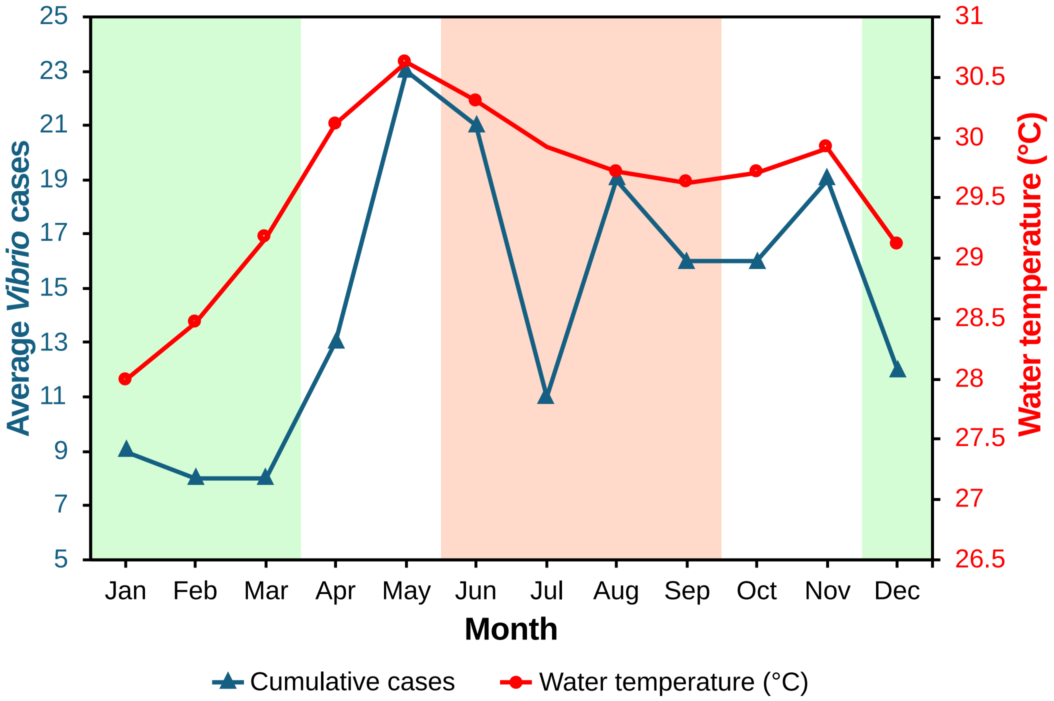


**Supplementary Fig. 5. Monthly variation of vibriosis cases versus water temperature in Singapore.** Vibriosis case totals and water temperature across the months of January to December. Shaded sections indicate the Southwest monsoon (June–September) and Northeast monsoon (December–March).

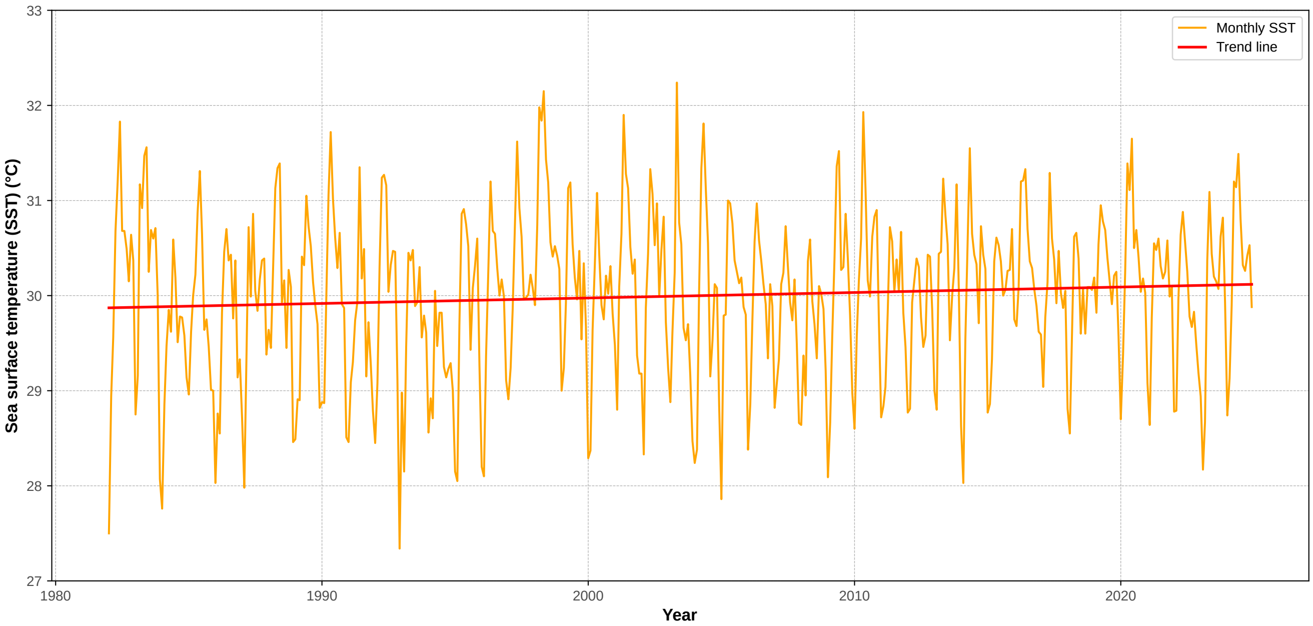


**Supplementary Fig. 6. Long-term trend in average sea surface temperature (SST) in Singapore (1982–2024).** Monthly SST near Singapore from NOAA's ¼° Daily OISST v2.1 monthly product, 1982–2024. The orange curve shows monthly SST; the red line represents the linear trend. The fitted slope is equivalent to ~0.058°C per decade, implying ~0.25°C warming over 1982–2024. Warmer coastal SSTs are associated in the literature with increased environmental suitability for marine pathogens and related infection risk.

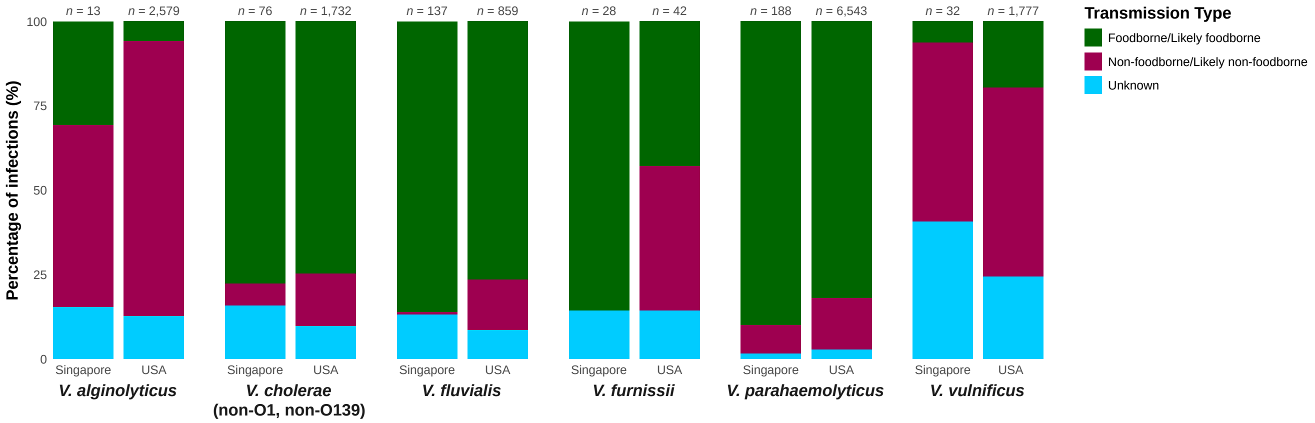


**Supplementary Fig. 7. Modes of transmission for vibriosis by species in the United States and Singapore.** Bar graphs show the proportion of infections attributed to different transmission pathways (i.e., foodborne/likely foodborne, non-foodborne/likely non-foodborne, and unknown) for each *Vibrio* species.

### SUPPLEMENTARY METHODS

##### Vibriosis case definitions

One consideration with vibriosis is that not all infections are symptomatic. For most *Vibrio* species, many infections are mild, self-limited illnesses that often go undiagnosed, with the exception of *Vibrio vulnificus*, which typically presents with more severe symptoms.[1] In the United States, reporting classifications are based on the Centers for Disease Control and Prevention (CDC) case definitions for probable and confirmed cases.[2] Currently, culture-independent diagnostic tests (CIDTs) are sufficient to meet the criteria for 'probable cases.' Culture confirmation is required for a case to be classified as 'confirmed,' which is the definition used in this study. If two or more *Vibrio* species are detected, each is considered a separate infection.[2] This approach was chosen to minimize the effects of CIDT introduction (in 2017) on apparent increases in diagnosis.

*Vibrio* species reported to the US COVIS database are based on results from state and local laboratories, which conduct culture confirmation using various methods such as API 20E or MALDI-TOF, both of which have limitations in sensitivity and specificity.[3] Whole genome sequencing-based identification was also available nationwide through PulseNet, the national molecular surveillance system for food- and waterborne pathogens, providing an additional layer of quality assurance for species identification.[4] For consistency in identification and reporting in this study, these CDC definitions were also applied to the Singapore dataset.

##### Statistical analysis

Monthly case counts were analyzed using a log-linked negative-binomial regression model to account for over-dispersion and to express incidence rates on a per capita basis.[5] The model included main effects for species and region, as well as their interaction, a linear time term (months since the first observation) and its interactions with both species and region, and an offset term of *log(Population)*. From this model, we obtained marginal time slopes (annual changes in incidence) for each species within each region; these were reported as rate ratios with Wald 95% confidence intervals. Resulting *p*-values were adjusted for multiple comparisons using the Benjamini–Hochberg false discovery rate procedure.[6]

### REFERENCES


1. Scallan, E. *et al.* Foodborne illness acquired in the United States—major pathogens. *Emerg Infect Dis* **17**, 7-15, doi:10.3201/eid1701.p11101 (2011).
2. US Centers for Disease Control and Prevention. *Vibriosis (any species of the family* Vibrionaceae*, other than toxigenic* Vibrio cholerae *O1 or O139) 2017 case definition*, <https://ndc.services.cdc.gov/case-definitions/vibriosis-2017> (April 16, 2021).
3. O'Hara, C. M., Sowers, E. G., Bopp, C. A., Duda, S. B. & Strockbine, N. A. Accuracy of six commercially available systems for identification of members of the family *Vibrionaceae*. *J Clin Microbiol* **41**, 5654-5659, doi:10.1128/JCM.41.12.5654-5659.2003 (2003).
4. Swaminathan, B., Barrett, T. J., Hunter, S. B., Tauxe, R. V. & Force, C. D. C. P. T. PulseNet: the molecular subtyping network for foodborne bacterial disease surveillance, United States. *Emerg Infect Dis* **7**, 382-389, doi:10.3201/eid0703.010303 (2001).
5. Linden, A. & Mäntyniemi, S. Using the negative binomial distribution to model overdispersion in ecological count data. *Ecology* **92**, 1414-1421, doi:10.1890/10-1831.1 (2011).
6. Benjamini, Y. & Hochberg, Y. Controlling the false discovery rate: a practical and powerful approach to multiple testing. *J R Stat Soc Series B Stat Methodol* **57**, 289-300, doi:10.1111/j.2517-6161.1995.tb02031.x (2018).
